# PredictProtein – Predicting Protein Structure and Function for 29 Years

**DOI:** 10.1101/2021.02.23.432527

**Authors:** Michael Bernhofer, Christian Dallago, Tim Karl, Venkata Satagopam, Michael Heinzinger, Maria Littmann, Tobias Olenyi, Jiajun Qiu, Konstantin Schütze, Guy Yachdav, Haim Ashkenazy, Nir Ben-Tal, Yana Bromberg, Tatyana Goldberg, Laszlo Kajan, Sean O’Donoghue, Chris Sander, Andrea Schafferhans, Avner Schlessinger, Gerrit Vriend, Milot Mirdita, Piotr Gawron, Wei Gu, Yohan Jarosz, Christophe Trefois, Martin Steinegger, Reinhard Schneider, Burkhard Rost

**Author notes:** The authors wish it to be known that, in their opinion, the first four authors should be regarded as joint First Authors. Corresponding author, http://rostlab.org, (email Rost).

## Abstract

Since 1992 *PredictProtein* (https://predictprotein.org) is a one-stop online resource for protein sequence analysis with its main site hosted at the Luxembourg Centre for Systems Biomedicine (LCSB) and queried monthly by over 3,000 users in 2020. *PredictProtein* was the first Internet server for protein predictions. It pioneered combining evolutionary information and machine learning. Given a protein sequence as input, the server outputs multiple sequence alignments, predictions of protein structure in 1D and 2D (secondary structure, solvent accessibility, transmembrane segments, disordered regions, protein flexibility, and disulfide bridges) and predictions of protein function (functional effects of sequence variation or point mutations, Gene Ontology (GO) terms, subcellular localization, and protein-, RNA-, and DNA binding). PredictProtein’s infrastructure has moved to the LCSB increasing throughput; the use of MMseqs2 sequence search reduced runtime five-fold; user interface elements improved usability, and new prediction methods were added. PredictProtein recently included predictions from deep learning embeddings (GO and secondary structure) and a method for the prediction of proteins and residues binding DNA, RNA, or other proteins. PredictProtein.org aspires to provide reliable predictions to computational and experimental biologists alike. All scripts and methods are freely available for offline execution in high-throughput settings.

**Availability:** *Freely accessible webserver* PredictProtein.org; Source and docker images: github.com/rostlab

## 1 Introduction

The sequence is known for far more proteins (1) than experimental annotations of function or structure (2, 3). This sequence-annotation gap existed when *PredictProtein* (4, 5) started in 1992, and has kept expanding ever since (6). Unannotated sequences contribute crucial evolutionary information to neural networks predicting secondary structure (7, 8) that seeded *PredictProtein (<u>PP</u>)* at the European Molecular Biology Laboratory (EMBL) in 1992 (9), the first fully automated, query-driven Internet server providing evolutionary information and structure prediction for any protein. Many other methods predicting aspects of protein function and structure have since joined under the PP roof (4, 5, 10) now hosted by the Luxembourg Centre of Systems Biomedicine (LCSB).

PP offers an array of structure and function predictions most of which combine machine learning with evolutionary information; now enhanced by a faster alignment algorithm (11, 12). A few prediction methods now also use embeddings (13, 14) from protein Language Models (LMs) (13–18). Embeddings are much faster to obtain than evolutionary information, yet for many tasks, perform almost as well, or even better (19, 20). All PP methods join at PredictProtein.org with interactive visualizations; for some methods, more advanced visualizations are linked (21–23). The reliability of *PredictProtein*, its speed, the continuous integration of well-validated, top methods, and its intuitive interface have attracted thousands of researchers over 29 years of steady operation.

## 2 Methods

### PredictProtein (PP) provides

multiple sequence alignments (MSAs) and position-specific scoring matrices (PSSMs) computed by a combination of pairwise BLAST (24), PSI-BLAST (25), and MMseqs2 (11, 12) on query vs. PDB (26) and query vs. UniProt (1). For each residue in the query, the following per-residue predictions are assembled: secondary structure (RePROF/PROFsec (5, 27) and ProtBertSec (14)); solvent accessibility (RePROF/PROFacc); transmembrane helices and strands (TMSEG (28) and PROFtmb (29)); protein disorder (Meta-Disorder (30)); backbone flexibility (relative B-values; PROFbval (31)); disulfide bridges (DISULFIND (32)); sequence conservation (ConSurf/ConSeq (33–36)); protein-protein, protein-DNA, and protein-RNA binding residues (ProNA2020 (3)); PROSITE motifs (37); effects of sequence variation (single amino acid variants, SAVs; SNAP2 (38)). For each query per-protein predictions include: transmembrane topology (TMSEG (28)); binary protein-(DNA|RNA|protein) binding (protein binds X or not; ProNA2020 (3)); Gene Ontology (GO) term predictions (goPredSim (19)); subcellular localization (LocTree3 (39)); Pfam (40) domain scans, and some biophysical features. Under the hood, PP computes more results (**Table S1**), either as input for frontend methods, or for legacy support.

### New: goPredSim embedding-based transfer of Gene Ontology (GO)

goPredSim (19) predicts GO terms by transferring annotations from the most embedding-similar protein. Embeddings are obtained from SeqVec (13); similarity is established through the Euclidean distance between the embedding of a query and a protein with experimental GO annotations. Replicating the conditions of CAFA3 (41) in 2017, goPredSim achieved F_max_ values of 37±2%, 52±2%, and 58±2% for BPO (biological process), MFO (molecular function), and CCO (cellular component), respectively (41, 42). Using annotations from the Gene Ontology Annotation (GOA) database (43, 44) in 2020 and testing on a set of 296 proteins with annotations added after February 2020 appeared to reach even slightly higher values that were confirmed through preliminary results for CAFA4 (45).

### New: ProtBertSec secondary structure prediction

ProtBertSec predicts secondary structure in three states (helix, strand, other) using Prot-Bert (14) embeddings derived from training on BFD with almost 3*10^9^ proteins (6, 46). On a hold-out set from CASP12, ProtBertSec reached a three-state per-residue accuracy of Q3=76±1.5% (47). Although below the state-of-the-art (e.g. NetSurfP 2 (48) at 82%), it appears to outperform all other existing methods while not using alignments.

### New: ProNA2020 predicts protein, RNA & DNA binding

ProNA2020 (3) predicts whether or not a protein interacts with other proteins, RNA or DNA, and if the binding residues. Per-protein predictions rely on homology and machine learning models employing profile-kernel SVMs (49) and embeddings from an *in-house* implementation of ProtVec (50). Per-residue predictions are based on simple neural networks due to the lack of experimental high-resolution annotations (51–53). ProNA2020 correctly predicted 77%±1% of proteins that bind DNA, RNA or protein. In proteins known to bind other proteins, RNA, or DNA, ProNA2020 correctly predicted 69±1%, 81±1%, and 80±1% of binding residues, respectively.

### New: MMseqs2 speedy evolutionary information

Most time-consuming for PP was the search for related proteins in ever growing databases. MMseqs2 (11) finds related sequences blazingly fast and seeds a PSI-BLAST search (25). The query sequence is sent to a dedicated MMseqs2 server that searches for hits against cluster representatives within the Uni-Clust30 (54) and PDB (26) reduced to 70% pairwise percentage sequence identity (PIDE). All hits and their respective cluster members are turned into a MSA and filtered to the 3,000 most diverse sequences.

## 3 Web Server

### Frontend and User Interface (UI)

Users query PredictProtein.org by submitting a protein sequence. Results are available in seconds for sequences that had been submitted recently (cf. *PPcache* next section), or within up to 30 minutes if predictions are recomputed. Per-residue predictions are displayed online via ProtVista (55), which allows to zoom into any sequential protein region (**Fig. S1**), and are grouped by category (e.g., secondary structure), which can be expanded to display more detail (e.g., helix, strand, other). On the results page, links to visualize MSAs through *AlignmentViewer* (56) are available. More predictions can be accessed through a menu on the left, e.g., *Gene Ontology Terms*, *Effect of Point Mutations* and *Subcellular Localization*. Prediction views include references and details of outputs, as well as rich visualizations, e.g., GO trees for GO predictions and cell images with highlighted predicted locations for subcellular localization predictions (57).

### PPcache, backend, and programmatic access

The PP backend lives at LCSB, allowing for up to 48 parallel queries. Results are stored on disc in the *PPcache* (5). Users submitting any of the 660,000 recently submitted sequences obtain results immediately. Using the bio-embeddings software (58), the PPcache is enriched by embeddings and embedding-based predictions on the fly. For all methods displayed on the frontend, JSON files compliant with *ProtVista* (55) are available via REST APIs (**SOM_1**), and are in use by external services such as the protein 3D structure visualization suite *Aquaria* (21, 23) and by *MolArt* (22).

### PredictProtein is available for local use

All results displayed on and downloadable from PP are available through Docker (59) and as source code for local and cloud execution (available at github.com/rostlab).

## 4 Use Case

We demonstrate PredictProtein.org tools through predictions of the novel coronavirus SARS-CoV-2 (NCBI:txid2697049) nucleoprotein (UniProt identifier P0DTC9/NCAP_SARS2; **Fig. 1**). NCAP_SARS2 has 419 residues and interacts with itself (experimentally verified). Sequence similarity and automatic assignment via UniRule suggest NCAP is RNA binding (binding with the viral genome), binding with the membrane protein M (UniProt identifier P0DTC5/VME1_SARS2), and is fundamental for virion assembly. goPredSim (19) transferred GO terms from other proteins for MFO (*RNA-binding*; GO:0003723; ECO:0000213) and CCO (compartments in the host cell and viral nucleocapsid; GO:0019013; GO:0044172; GO:0044177; GO:0044220; GO:0030430; ECO:0000255) matching annotations found in UniProt (1). While it missed the experimentally verified MFO term *identical protein binding* (GO:0042802), goPredSim predicted *protein folding* (GO:0006457) and *protein ubiquitination* (GO:0016567) suggesting the nucleoprotein to be involved in biological processes requiring protein binding. ProNA2020 (3) predicts RNA-binding regions, the one with highest confidence between I84 (Isoleucine at position 84) and D98 (Aspartic Acid at 98) (protein sequence available in **SOM_1**). While high resolution experimental data on binding is not available, an NMR structure of the SARS-CoV-2 nucleocapsid phospho-protein N-terminal domain in complex with 10mer ssRNA (PDB identifier 7ACT (60)) supports the predicted RNA-binding site (**Fig. 2**). Additionally, SNAP2 (38) predicts single amino acid variants (SAVs) in that region to likely affect function, reinforcing the hypothesis that this is a functionally relevant site. Although using different input information (evolutionary vs. embeddings), RePROF (5) and ProtBertSec (14) both predict an unusual content exceeding 70% non-regular (neither helix nor strand) secondary structure, suggesting that most of the nucleoprotein might not form regular structure. This is supported by Meta-Disorder (30) predicting 53% overall disorder. Secondary structure predictions match well high-resolution experimental structures of the nucleoprotein not in complex (e.g., PDB 6VYO (61); 6WJI (62)). Both secondary structure prediction methods managed to zoom into the ordered regions of the protein and predicted e.g., the five beta-strands that are formed within the sequence range I84 (Isoleucine) to A134 (Alanine), and the two helices formed within the sequence range spanned from F346 (Phenylalanine) to T362 (Tyrosine).

**Fig. 1:**
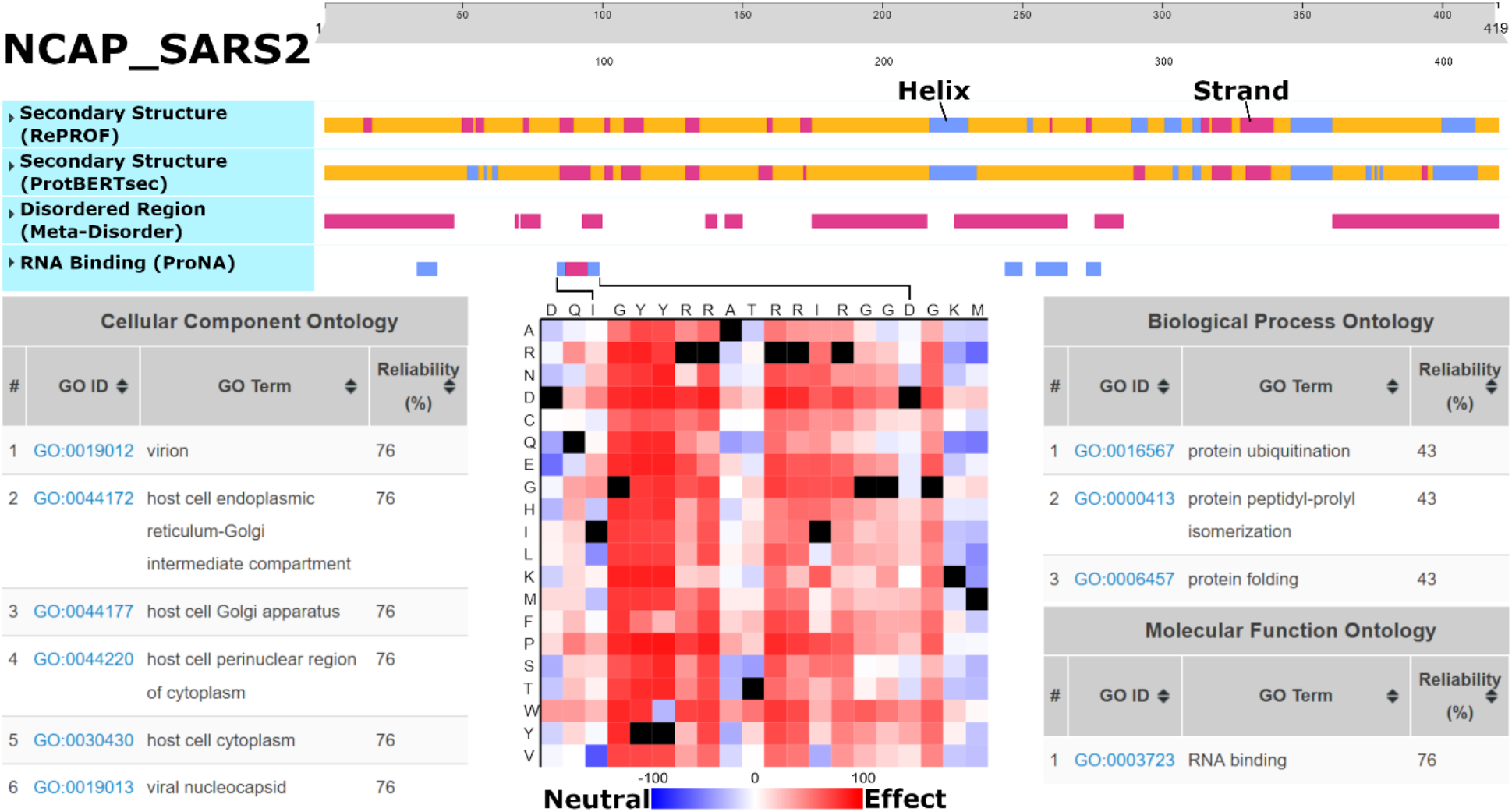
Predictions for SARS-CoV-2 Nucleoprotein (NCAP_SARS2). Underneath the interactive slider at the top: RePROF and ProtBertSec secondary structure (blue: helix; purple: strand; orange: other); Meta-Disorder intrinsically disordered regions (purple); ProNA2020 RNA-binding residues (low confidence: blue; medium confidence: purple). goPredSim transfers of GeneOntology (GO) terms based on embedding similarity (lower left: CCO; lower right: BPO & MFO). SNAP2 predicts the effect on function of point-mutations for the RNA binding region from I84 to D98 (bottom-center; black: native residue). Link: predictprotein.org/visual_results?req_id=$1$nAmulUQY$FRPFaP8NTqLW9DzdlTG3B/

**Fig. 2:**
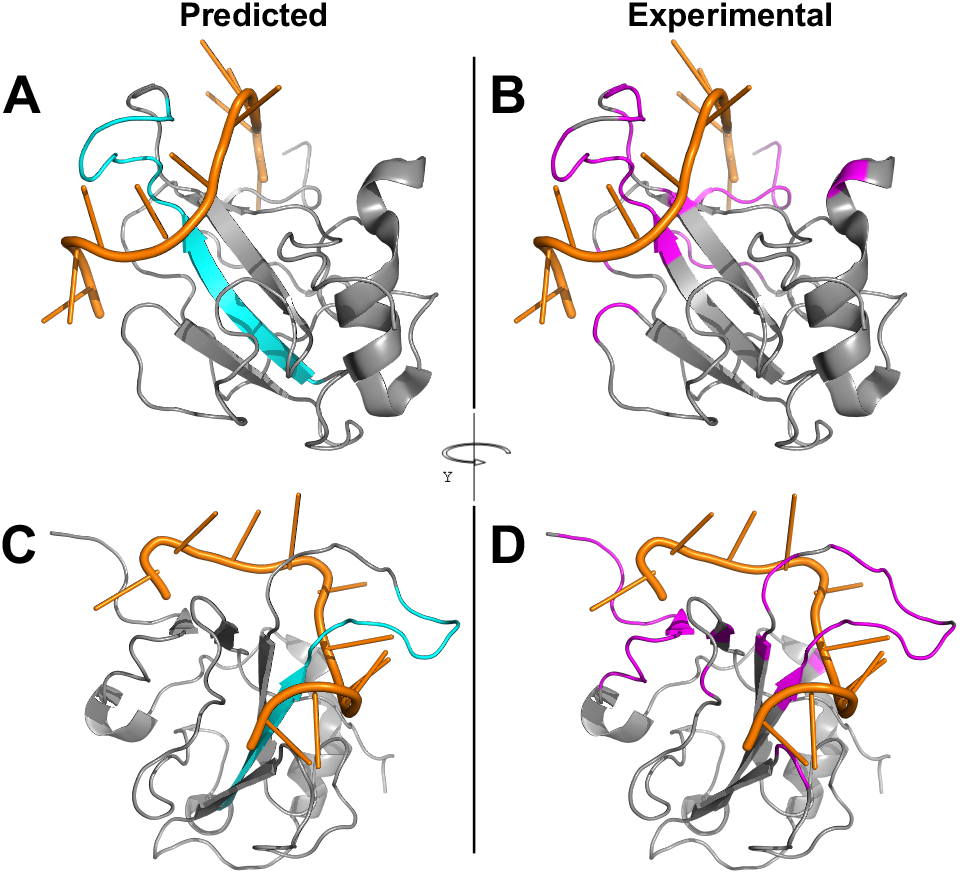
Experimental and predicted RNA binding residues for NCAP2_SARS2. Predicted (via ProNA2020, in cyan, Panels A and C) and observed (within 5 Ångstrøm, in magenta, Panels B and D) RNA binding residues for the SARS-cov-2 nucleoprotein (gray) complexed with a 10-mer ssRNA (orange), PDB structure 7ACT (60). Two-third of the predictions are correct (precision=0.73, recall=0.20), which is around the expected average performance reported by ProNA2020. The important sequence consecutive central strand and loop are predicted well, while several short sequence segments that are far away in sequence space but close in structure space are missed, which is expected as ProNA2020 has no notion of 3D structure, i.e., cannot identify “binding sites”. A-B show a different orientation than C-D.

## 5 Conclusion

For almost three decades (preceding Google) *PredictProtein* (PP) offers predictions for proteins. PP is the oldest Internet server in protein prediction, online for 6-times as long as most other servers (63–65). It pioneered combining machine learning with evolutionary information and making predictions freely accessible online. While the sequence-annotation gap continues to grow, the sequence-structure gap might be bridged in the near future (66). For the time being, servers such as PP, providing a one-stop solution to predictions from many sustained, novel tools are needed. PP is the first server to offer fast embedding-based predictions of structure (ProtBertSec) and function (goPredSim). By slashing runtime for PSSMs from 72 to 4 minutes through MMseqs2 and better LCSB hardware, PP also delivers evolutionary information-based predictions fast! Instantaneously if the query is in the precomputed *PPcache*. For heavy use, the of-fline Docker containers provide predictors out-of-the-box. At no cost to users, *PredictProtein* offers to quickly shine light on proteins for which little is known using well validated prediction methods.

## Supporting information

SOM

## Acknowledgements

Maintaining *PredictProtein* over three decades has been tough; many colleagues have helped with hands and brains, developers, and users alike. Thanks to all of you! Please find most contributors in **Table S2** or at predictprotein.org/credits. In particular, thanks to Noua Toukourou and Maharshi Vyas (both LCSB) for invaluable help with hardware and software; to David Hoksza (Charles U, Prague) for his work on MolArt; to Marco Punta (IRCCS Milano) for his long-term support; to Inga Weise (TUM) for support with many aspects; to Roy Omond (Blue Bubble, Cambridge), Antoine de Daruvar (Univ. Bordeaux), Yanay Ofran (Bar-Ilan Univ.), Jinfeng Liu (Genentech), Tobias Hamp, Maximilian Hecht, Edda Kloppmann (all previously TUM) for contributing methods and code in the past; Johannes Söding for providing resources to develop and maintain MMseqs2.

