## Supplementary material for "PredictProtein – Predicting Protein Structure and Function for 29 Years": SOM

#### Table of Contents for Supporting Online Material

### Additional Figures and Tables

#### Fig. S1: ProtVista Feature Viewer in PredictProtein

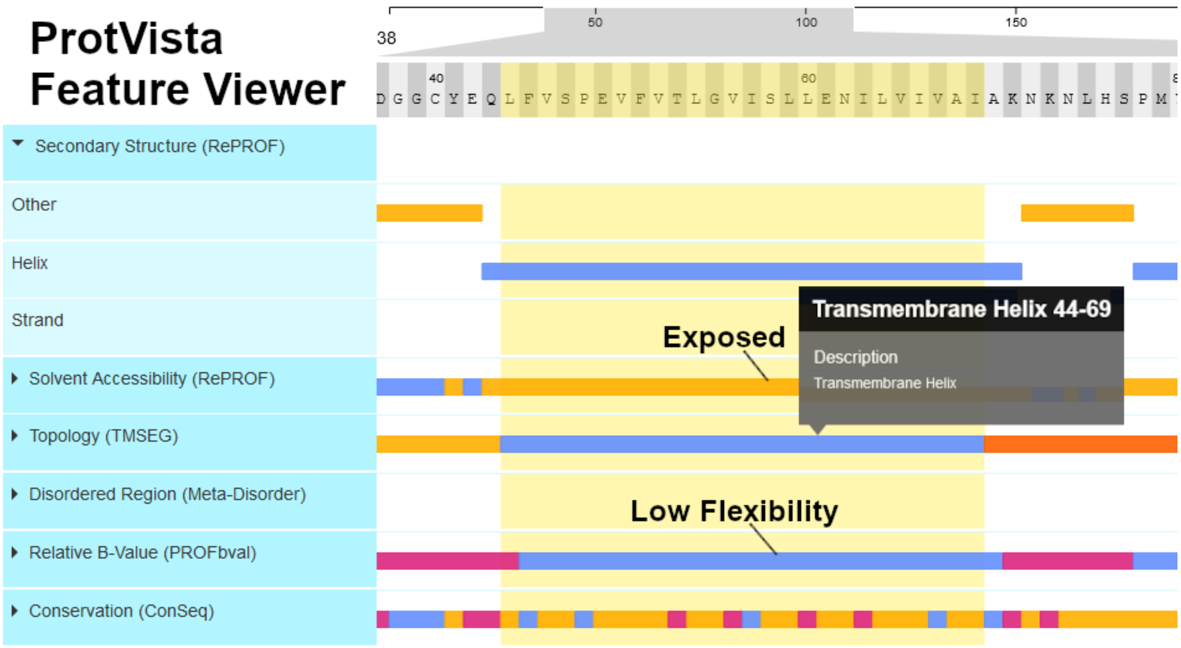

**Fig. S1: ProtVista Feature Viewer in PredictProtein.** The ProtVista (1) feature viewer allows to zoom into sequence regions (indicated by the grey range at the top). Predicted feature tracks can be expanded into sub-tracks (e.g., secondary structure: other, helix, strand). Clicking a segment (coloured bar) in the viewer prompts a tooltip details and highlights the region in the viewer in yellow (here: transmembrane helix from position 44 to 69). On the PredictProtein dashboard, the yellow highlight enables comparison of predicted features. The predicted transmembrane helix (TMH) matches with the predicted secondary structure (helix), solvent accessibility (exposed) and low flexibility within this region. The TMH is highly conserved as indicated by ConSeq (yellow: high conservation).
Link: predictprotein.org/visual_results?req_id=$1$gqwU35tl$5PIDUrsPtT7S57wVQZDCf0

#### Table S1: List of PredictProtein methods

Table of all methods executed by PredictProtein and brief description of their individual tasks. Availability provides information about whether the results are visualized on the website, available for download as binary files, and if so, their extension (all files share the same base name: “query”, e.g. “query.chk”). Predictions marked with “Web” can be downloaded as binary files from the PredictProtein frontend. SNAP2 and embedding-based predictions are not stored in the PPcache but are available for download via REST calls (see next section).

| **Method** | **Task** | **Availability** | | |
|---|---|---|---|---|
| | | Visual | Download | File(s) |
| BLAST (2), PSI-BLAST (3) | Pairwise alignments, PSSM generation | X | X | blastPsiAli.gz, blastPsiMat, blastPsiRdb, blastpSwissM8, chk |
| MMseqs2 (4) | Pairwise alignments | X | X | mmseqs2AliPdb, mmseqs2AliUref |
| ConSeq (5) | Amino acid conservation | X | X | consurf.grades, consurf.html |
| RePROF | Prediction of secondary structure and solvent accessibility | X | X | reprof |
| TMSEG (6) | Prediction of transmembrane helices | X | X | tmseg |
| Meta-Disorder (7) | Consensus prediction of disordered regions | X | X | mdisorder |
| PROFbval (8) | Prediction of residue mobility | X | X | profbval, profb4snap |
| DISULFIND (9) | Prediction of disulfide bridges | X | X | disulfinder |
| LocTree3 (10) | Prediction of sub-cellular localization for all domains of life | X | X | arch.lc3, arch.lc3.pb, arch.lc3.svm, bact.lc3, bact.lc3.pb, bact.lc3.svm, euka.lc3, euka.lc3.pb, euka.lc3.svm |
| ProNA2020 (11) | Prediction of protein-protein, -DNA and -RNA binding proteins and sites | X | X | prona |

| | | | | |
|---|---|---|---|---|
| SNAP2 (12) | Prediction of functional changes due to single amino acid variants | X | Web | |
| ProtBert-BFD-sec (13) | Prediction of secondary structure (embeddings input) | X | Web | |
| goPredSim (14) | Prediction of GO terms (embeddings input) | X | Web | |
| Metastudent (15) | Prediction of GO terms | | X | metastudent.BPO.txt, metastudent.CCO.txt, metastudent.MFO.txt |
| NORSnet (16) | Prediction of disordered regions | | X | norsnet |
| Norsp (17) | Prediction of non-regular secondary structure | | X | nors, sumNors |
| TMHMM (18) | Prediction of transmembrane helices | | X | tmhmm |
| PHDhtm (19) | Prediction of transmembrane helices | | X | phdPred, phdRdb |
| PROFtmb (20) | Prediction of transmembrane beta-barrels | | X | proftmb, proftmbdat |
| PROFacc (21), PROFsec (22) | Prediction of secondary structure and solvent accessibility | | X | profRdb, prof1Rdb, profAscii |
| PROFisis (23) | Prediction of protein-protein binding sites | | X | isis |
| PROFdisis (24) | Prediction of protein-DNA binding sites | | X | disis |
| SomeNA | Prediction of protein-DNA and -RNA binding sites | | X | somena |
| COILS (25) | Prediction of coiled coils | | X | coils, coils_raw |

| | | | | |
|---|---|---|---|---|
| PredictNLS (26) | Prediction of nuclear localization signals | | X | nls,<br>nlsDat,<br>nlsSum |
| SEG (27) | Mask low complexity regions | | X | segNorm |
| HMMER (28) | Search query for Pfam (29) domains | | X | hmm2pfam,<br>hmm3pfam,<br>hmm3pfamTbl,<br>hmm3pfamDomTbl |
| PROSITE (30) | Scan query for PROSITE patterns | | X | prosite |
| PSIC (31) | Profile extraction from alignments | | X | psic,<br>clustalngz |
| HSSP | Homology derived secondary structure | | X | hsspPsiFil.gz |

**Table S2: List of contributors**

List of all non-coauthor contributors. All contributors are acknowledged at https://predictprotein.org/credits

| **Name** | **Contribution** |
|---|---|
| Henry Bigelow | Contributed the PROFtmb method |
| Juan Miguel Cejuela | Added the literature search feature |
| Antoine de Daruvar | Helped getting the first PredictProtein server online |
| Rachel First | Designed the artwork for the localization prediction<br>Desinged the Site Tutorial |
| Paolo Frasconi | Contributed the DISULFIND method |
| Tobias Hamp | Contributed the Metastudent method |
| Maximilian Hecht | Contributed the SNAP2 method |
| David Hoksza | Developer of MolArt |
| Peter Hönigschmid | Contributed the SomeNA method |
| Edda Kloppmann | Scientific advisor |
| Jinfeng Liu | Contributed code for the PredictProtein pipeline<br>Contributed the NORS, CHOP & CHOPnet methods |

| | |
|---|---|
| Sven Mika | Contributed the UniqueProt method |
| Rajesh Nair | Contributed the LocTree method |
| Yanay Ofran | Contributed the PPSites and PROFdisis methods |
| Roy Omond | Helped in the communication between VMS and Unix systems for the first server |
| Dariusz Przybylski | Contributed the AGAPE metod |
| Marco Punta | Contributed the Meta-Disorder method |
| Jonas Reeb | Scientific advisor |
| Lothar Richter | Scientific advisor |
| Manfred Roos | Maintained the PredictProtein Knowledgebase |
| Thomas Splettstoesser | Designed the PredictProtein logo |
| Noua Toukourou | Hosts and supports the PredictProtein server at the LCSB |
| Maharshi Vyas | Hosts and supports the PredictProtein server at the LCSB |
| Kazimierz Wrzeszczynski | Contributed code and ideas |

**Table S3: goPredSim predictions for NCAP_SARS2**

GO terms inferred by goPredSim for the SARS-COV-2 nucleoprotein NCAP_SARS2.

| ***Cellular Component Ontology*** | | |
|---|---|---|
| **GO Term** | **GO ID** | **Reliability (%)** |
| virion | GO:0019012 | 76 |
| host cell endoplasmic reticulum-Golgi intermediate compartment | GO:0044172 | 76 |
| host cell Golgi apparatus | GO:0044177 | 76 |
| host cell perinuclear region of cytoplasm | GO:0044220 | 76 |
| host cell cytoplasm | GO:0030430 | 76 |
| viral nucleocapsid | GO:0019013 | 76 |
| ***Biological Process Ontology*** | | |

| GO Term | GO ID | Reliability (%) |
|---|---|---|
| protein ubiquitination | GO:0016567 | 43 |
| protein peptidyl-prolyl isomerization | GO:0000413 | 43 |
| protein folding | GO:0006457 | 43 |
| ***Molecular Function Ontology*** | | |
| **GO Term** | **GO ID** | **Reliability (%)** |
| RNA binding | GO:0003723 | 76 |

### Programmatic access via PredictProtein APIs

Essentially, the PredictProtein web server has two APIs: one to access the result files of the main PredictProtein pipeline and prediction methods (stored in the *PPCache*), and one for the ProtVista viewer formatted JSON output. Further, there are two more APIs for the external web services of BioEmbeddings and SNAP2.

***PredictProtein File API.*** Result files generated by PredictProtein and stored in the PPCache can be queried and downloaded directly via API (in addition to the web interface). The API can be accessed via POST requests with a JSON payload to https://predictprotein.org/api/ppc_fetch. The request must include the type of action to be performed (*get* or *has*) and the protein sequence. Optionally, it can include one or more method names (comma separated), or a specific file name (but not methods and file name). A *has*-request returns the available files in the PPCache for the requested sequence. If one or more methods or a file name has been specified, the returned file list will only include the corresponding files, if available. A *get*-request returns all requested and available files as a zip archive. If only a single file is requested (via file name), the raw unzipped file is returned instead.

List of supported methods: *coils, conseq, disulfind, hmmer, hssp, loctree3, mdisorder, mmseqs2, mstudent, norsnet, norsp, phdhtm, predictnls, profacc, profbval, profdisis, profisis, profsec, proftmb, prona, prosite_scan, psiblast, psic, reprof, seg, somena, tmhmm, tmseg.*

All files listed for the different methods (Table S1) are supported by the API. Each file starts with the same base name "query" and has the listed file extension. For example, the result file for TMSEG is: *query.tmseg*

Example POST requests (JSON payload) for sequence "SEQWENCE":

- Get list of all available files:
  *{"action": "has", "sequence": "SEQWENCE"}*
- Download all files for LocTree3 and RePROF:
  *{"action": "get", "sequence": "SEQWENCE", "method": "loctree3,reprof"}*
- Download the file query.tmseg:
  *{"action": "get", "sequence": "SEQWENCE", "file": "query.tmseg"}*

***PredictProtein ProtVista API.*** Prediction results for a limited list of methods can be exported in JSON format compatible with the ProtVista viewer. The API can be accessed via POST requests with a JSON payload including the protein sequence to https://api.predictprotein.org/v1/results. The returned JSON output includes all available results for the following methods: consurf, disulfind, mdisorder, norsnet, profbval, prona, reprof, tmseg.

Example POST request (JSON payload) for sequence “SEQWENCE”:

- Get ProtVista JSON output:
  *{"protein": {"sequence": "SEQWENCE"}}*

***SNAP2 API***. SNAP2 predictions are available in JSON format via GET requests to https://rostlab.org/services/aquaria/snap4aquaria/json.php. The GET request must include the protein sequence. Using the optional *details* parameter, the JSON output can be switched between a summary report for each position, or a detailed list of all substitutions at every position.

Example GET request for sequence “SEQWENCE”:

- Get SNAP2 summary:
  https://rostlab.org/services/aquaria/snap4aquaria/json.php?seq=SEQWENCE
- Get detailed SNAP2 report:
  https://rostlab.org/services/aquaria/snap4aquaria/json.php?details&seq=SEQWENCE

***BioEmbeddings API***. Embedding-input-based predictions are available in various JSON formats via POST and GET requests using a sequence directly. Detailed documentation is available at https://embeddings.predictprotein.org/api.

Example POST request (JSON payload) for sequence “SEQWENCE”:

POST to https://embeddings.predictprotein.org/api/annotations with JSON payload:
*{"model": “seqvec”, "sequence": "SEQWENCE", “format”: “go-predictprotein”}*

### Sequence information

##### Protein sequence of SARS-CoV-2 (NCBI:txid2697049) nucleoprotein (UniProt identifier P0DTC9/ncap_sars2)

```
>sp|P0DTC9|NCAP_SARS2 Nucleoprotein OS=Severe acute respiratory syndrome coronavirus 2 OX=2697049 GN=N PE=1 SV=1
MSDNGPQNQRNAPRITFGGPSDSTGSNQNGERSGARSKQRRPQGLPNNTASWFTALTQHG
KEDLKFPRGQGVPINTNSSPDDQIGYYRRATRRIRGGDGKMKDLSPRWYFYYLGTGPEAG
LPYGANKDGIIWVATEGALNTPKDHIGTRNPANNAAIVLQLPQGTTLPKGFYAEGSRGGS
QASSRSSSRSRNSSRNSTPGSSRGTSPARMAGNGGDAALALLLLDRLNQLESKMSGKGQQ
QQGQTVTKKSAAEASKKPRQKRTATKAYNVTQAFGRRGPEQTQGNFGDQELIRQGTDYKH
WPQIAQFAPSASAFFGMSRIGMEVTPSGTWLTYTGAIKLDDKDPNFKDQVILLNKHIDAY
KTFPPTEPKKDKKKKADETQALPQRQKKQQTVTLLPAADLDDFSKQLQQSMSSADSTQA
```
